# Tau deposition is associated with imaging patterns of tissue calcification in the P301L mouse model of human tauopathy

**DOI:** 10.1101/851915

**Authors:** Ruiqing Ni, Yvette Zarb, Gisela A. Kuhn, Ralph Müller, Yankey Yundung, Roger M. Nitsch, Luka Kulic, Annika Keller, Jan Klohs

## Abstract

Brain calcification is associated with several neurodegenerative proteinopathies. Here, we report a new phenotype of intracranial calcification in transgenic P301L mice overexpressing 4 repeat tau. P301L mice (Thy1.2) of 3, 5, 9 and 18-25 months-of-age and age-matched non-transgenic littermates were assessed using *in vivo/ex vivo* magnetic resonance imaging (MRI) with a gradient recalled echo sequence and micro computed tomography (μCT). Susceptibility weighted images computed from the gradient recalled echo data revealed regional hypointensities in the hippocampus, cortex, caudate nucleus and thalamus of P301L mice, which in corresponding phase images indicated diamagnetic lesions. Concomitantly, µCT detected hyperdense lesions. Occurrence of diamagnetic susceptibility lesions in the hippocampus, increased with age. Immunochemical staining of brain sections revealed bone protein-positive deposits. Furthermore, intra-neuronal and vessel-associated protein-containing nodules co-localized with phosphorylated-tau (AT8 and AT100) in the hippocampus. Protein-containing nodules were detected also in the thalamus in the absence of phosphorylated-tau deposition. In contrast, osteocalcin-containing nodules were vessel-associated, indicating ossified vessels, in the thalamus in absence of phosphorylated-tau. In summary, MRI and µCT demonstrated imaging pattern of intracranial calcification, concomitant with immunohistochemical evidence of formation of protein deposits containing bone proteins along with phosphorylated-tau in the P301L mouse model of human tauopathy. The P301L mouse model may thus serve as a future model to study the pathogenesis of brain calcifications in tauopathies.

## Introduction

Brain calcifications are detected in approximately 20% of aged individuals undergoing diagnostic neuroimaging, particularly in the pineal gland and choroid plexus [9, 15, 20, 29]. Higher incidences of brain calcifications are found in patients with neurodegenerative diseases such as Alzheimer’s disease (AD) [37, 38], cerebral amyloid angiopathy [4], frontotemporal dementia (FTD) [36], Parkinson’s disease and Down syndrome [68]. Calcified deposits in the human brain are also found distributed throughout the hippocampus and basal ganglia, and some are associated with cerebral vessels and cerebral amyloid angiopathy [7, 9, 15, 20, 57]. In patients with diffuse neurofibrillary tangles with calcification, pallidal calcification is the most prominent feature and is accompanied by frontotemporal atrophy, neurofibrillary tangle (NFT) deposition throughout the neocortex, without the occurrence of β-amyloid plaques [30]. Symmetrical calcification of the basal ganglia is a key diagnostic criterion in patients with primary familial brain calcification [70]. The etiology of brain calcification is poorly understood, despite high prevalence of calcification in neurodegenerative disorders.

The discovery of tau mutations has facilitated the generation of several mouse models of human tauopathy e.g. P301S and P301L lines (transgenic for a human 4 repeat tau isoform) [6, 52–54], which have become important tools to study the mechanisms of abnormal tau aggregation and deposition in FTD (4 repeat tau) [61] and AD (3 and 4 repeat tau) [25, 33, 41]. In the P301L (Thy 1.2, pR5 line) [11, 18], P301L (CaMKIIa) [23, 43, 62] and P301L (tetO) [14] mouse models, tau deposits begin forming before 3 months-of-age in neurons in the entorhinal cortex, hippocampus and later in the cortex, and amygdala; with neuroinflammation and impaired memory functions in hippocampus- and amygdala-dependent tasks manifesting at a later stage [49, 50, 67]. Brain atrophy and white matter changes indicating neurodegeneration were reported in P301L (CaMKIIa) mouse line around 9 months-of-age [16, 23, 43, 62], which was driven by factors additional to human tau overexpression. However, unlike in human patients with tauopathies, brain calcifications have not been reported in transgenic mouse models of human disease.

Here, we report an early occurrence of intracranial calcifications in the P301L (Thy1.2) mouse model of tauopathy. Evidence is provided by high-field magnetic resonance imaging (MRI), micro computed tomography (μCT) which correspond to protein-containing nodules upon immunohistochemical staining. Parenchymal and vascular calcifications increase with age and are associated with intra-neuronal phosphorylated-tau deposition in the hippocampus. The P301L mouse line may be a suitable animal model to study the role of brain calcification in human tauopathy.

## Materials and methods

### Animals

Homozygous mice, transgenic for a human four repeat isoform with the P301L under Thy1.2 promoter (C57B6.Dg background) [19] and non-transgenic littermates were used (see groups in **Supplementary Table 1**). Animals were housed in individually ventilated cages inside a temperature-controlled room, under a 12-hour dark/light cycle. Pelleted food (3437PXL15, CARGILL) and water was provided *ad-libitum*. All experiments were performed in accordance with the Swiss Federal Act on Animal Protection and approved by the Cantonal Veterinary Office Zurich (permit number: ZH082/18).

### Magnetic resonance imaging

MRI was performed as described previously [27, 44]. *In vivo* MRI was performed at 7T for detecting calcifications and volumetry. A comparison between the two field strengths was performed in order to determine if using a lower magnetic field at 7T (feasible in a clinical setting), could detect the diamagnetic signal from calcification. *In vivo* MRI was completed on a Bruker Biospec 70/40 (Bruker Biospin GmbH, Ettlingen, Germany) small animal MR system equipped with an actively shielded gradient set of 760 mT/m and 80 μs rise time and operated by a Paravision 6.0.1 software platform (Bruker Biospin GmbH, Ettlingen, Germany). Mice were anesthetized with an initial dose of 4 % isoflurane in oxygen/air (200:800 ml/min) and maintained at 1.5 % isoflurane in oxygen/air (100:400 ml/min). Body temperature was monitored with a rectal temperature probe (MLT 415, AD Instruments, Spechbach, Germany) and kept at 36.5 ± 0.5 °C on a water-heated holder (Bruker BioSpin AG, Fällanden, Switzerland). For susceptibility weighted imaging (SWI), a two-dimensional flow compensated gradient-recalled echo (FLASH) sequence was applied with the following parameters: field-of-view = 20×20 mm; image size = 256×256 mm, slice thicknesss = 0.8 mm, resulting in a resolution of 78×78 µm, number of slices = 20. One echo with an echo time = 18 ms; repetition time = 698 ms; flip angle= 30 °; and number of averages = 30 within an acquisition scan time of 1 h 29 min 2 s was recorded. Global 1^st^-order shimming followed by fieldmap-based local shimming on the mouse brain was performed using the automated MAPshim routine, with an ellipsoid reference volume covering the whole cerebrum.

To compare the image quality of data acquired *in vivo* at 7T, *ex vivo* MRI was done on a Bruker Biospec 94/30 (Bruker Biospin GmbH, Ettlingen, Germany) small animal MR system. The system was equipped with a cryogenic 2×2 radiofrequency surface coil probe (Bruker BioSpin AG, Fällanden, Switzerland). After *in vivo* MRI, mice were perfused under ketamine/xylazine/acepromazine maleate anesthesia (75/10/2 mg/kg body weight, i.p. bolus injection) with 0.1 M PBS (pH 7.4) and decapitated. The mouse heads were post-fixed in 4 % paraformaldehyde in 0.1 M PBS (pH 7.4) for 6 days and stored in 0.1 M PBS (pH 7.4) at 4 °C afterwards. The heads were then placed in a 15 ml centrifuge tube filled with perfluoropolyether (Fomblin Y, LVAC 16/6, average molecular weight 2700, Sigma-Aldrich, U.S.A.). Samples were measured at room temperature. The brains with skull were scanned to detect calcification using SWI sequence. A 3D gradient-recalled echo SWI sequence was recorded with the following parameters: field-of-view = 15×12×15 mm; image size = 248×200×36 mm, resulting in a spatial resolution of 60×60×417 µm. One echo with an echo time = 12 ms; repetition time = 250 ms; flip angle= 15 °; number of averages = 4 within an acquisition scan time of 1 h 59 min 24 s was recorded. To reduce field inhomogeneities, global 1^st^-order followed by fieldmap-based local shimming on the mouse brain was performed. Phase maps and SW images were generated using Paravision software 6.0.1 as described previously [27].

SW images were compared with their phase image counterparts to ensure that the signals were due to a diamagnetic signal (i.e. presence of calcifications). The numbers of suspected calcified spots in a panel of an anatomical region (hippocampus, thalamus, caudate nucleus, midbrain and cortex etc.) were quantified using the *ex vivo* datasets acquired at 9.4T. The Allen mouse brain atlas was used for anatomical reference [24].

### µCT

After the *ex vivo* MRI, the mouse heads were scanned in 0.1 M PBS (pH 7.4) in a microCT 40 (Scanco Medical AG, Brüttisellen, Switzerland) operated at 45 kVp, 177 µA intensity, an integration of 200 ms and two-fold frame averaging. From 1000 projection images, 3D datasets with isotropic voxels of 8µm were reconstructed and converted to DICOM format using the scanner software. DICOM files were exported and analyzed using ITK SNAP [73].

### Histochemistry & Immunohistochemistry

The 4 % paraformaldehyde fixed mouse brains were cut sagittally into two hemispheres. One hemisphere was cut into 60 μm coronal sections using a vibratome (Leica VT1000S, Germany) for fluorescence immunohistochemistry. The protocol used for fluorescence immunohistochemistry was described previously [43, 46, 75]. Primary antibodies used for immunofluorescence staining are listed in **Supplementary table 2**. All fluorescently-labelled (Alexa 488, Cy3, DyLight 649) secondary antibodies (suitable for multiple labelling) were hosted in donkey (anti-rabbit, anti-rat, anti-mouse and anti-goat, Jackson Immunoresearch, UK). Brain sections were pre-treated with mouse-on-mouse kit (Vector Laboratories, USA) to quench the endogenous IgG and subsequently incubated with primary and secondary antibodies. Immunohistochemistry stainings were imaged using a confocal microscope (Leica SP5; 40 × numerical aperture: 1.25; 63 × numerical aperture: 1.4). Images were analyzed using the image-processing software Imaris 8.4.1 (Bitplane, USA) and Illustrator CS 6 (Adobe, USA).

For histochemistry and non-fluorescent immunostaining, the other brain hemisphere was embedded in paraffin following routine procedures and cut into 2 μm thick sections. Sections were stained using Hematoxylin & Eosin, Alcian blue, Periodic acid–Schiff or Prussian blue using a standard protocol. The sections were deparaffinized and rehydrated before immunostaining. For glial fibril acidic protein (GFAP) staining, sections were incubated with rabbit anti-GFAP (DakoCytomation A/S, Denmark, #20334), followed by an incubation with HRP-conjugated goat anti-rabbit (Jackson Immunoresearch, USA). For ionized calcium binding adaptor molecule 1 (Iba1) staining, antigen retrieval was performed using hot citrate buffer (0.01 M; pH 6), followed by an incubation with rabbit anti-Iba1 (WAKO, Japan; 1:2500) and subsequently a biotinylated secondary antibody (Vector laboratories, USA). Visualization ensued after using the ABC complex solution (Vector laboratories, USA), 3,3′-Diaminobenzidine (DAB; Sigma-Aldrich, Switzerland) and hydrogen peroxide (Sigma-Aldrich, Switzerland). Counterstaining was performed using Hematoxylin. Stained paraffin sections were scanned using a NanoZoomer HT (Hamamatsu Photonics, Japan) using a 40× objective. Images were analyzed using Digital Image Hub software (SlidePath) and Adobe Illustrator CS6 (Adobe, USA). Hematoxylin & Eosin stained whole sagittal mouse brain slices were imaged at 20× magnification using Pannoramic 250 (3D HISTECH, Hungary). The images were analyzed using CaseViewer (3D HISTECH, Hungary) and ImageJ (NIH, USA).

### Statistics

Statistical analysis was performed using GraphPad Prism 7.0 (GraphPad Software, U.S.A). D’Agostino & Person normality test was used for assessing the normal distribution of the data. One-way ANOVA with Turkey’s post-hoc analysis was used for group comparison. The difference between groups was considered significant (*) at *p* value < 0.05. All error bars in the figures are expressed as standard deviation.

## Results

### Phase imaging detects the presence of diamagnetic lesions in P301L mouse brain

Gradient recalled echo data of P301L mice and non-transgenic littermates were collected (**Fig. 1-3**). We observed hypointensities in the SW images (**Fig. 1-3**). Corresponding phase images showed positive phase shifts (hyperintensities) in corresponding locations, indicating the diamagnetic nature of lesions (**Fig. 1-3**). The magnetic susceptibility [55] and appearance of the lesions indicated the presence of calcified deposits. Blood degradation products such as hemosiderin are paramagnetic and would induce opposite signal shifts. Hypointensities in the SW images on reconstructed data from *in vivo* MRI at 7T (**Fig. 1**) correspond well to that from *ex vivo* MRI at 9.4T (**Fig. 3**). However, boundaries of lesions were sharper in the 9.4T compared to 7T images due to the difference in field strengths and scan parameters.

**Figure 1.**
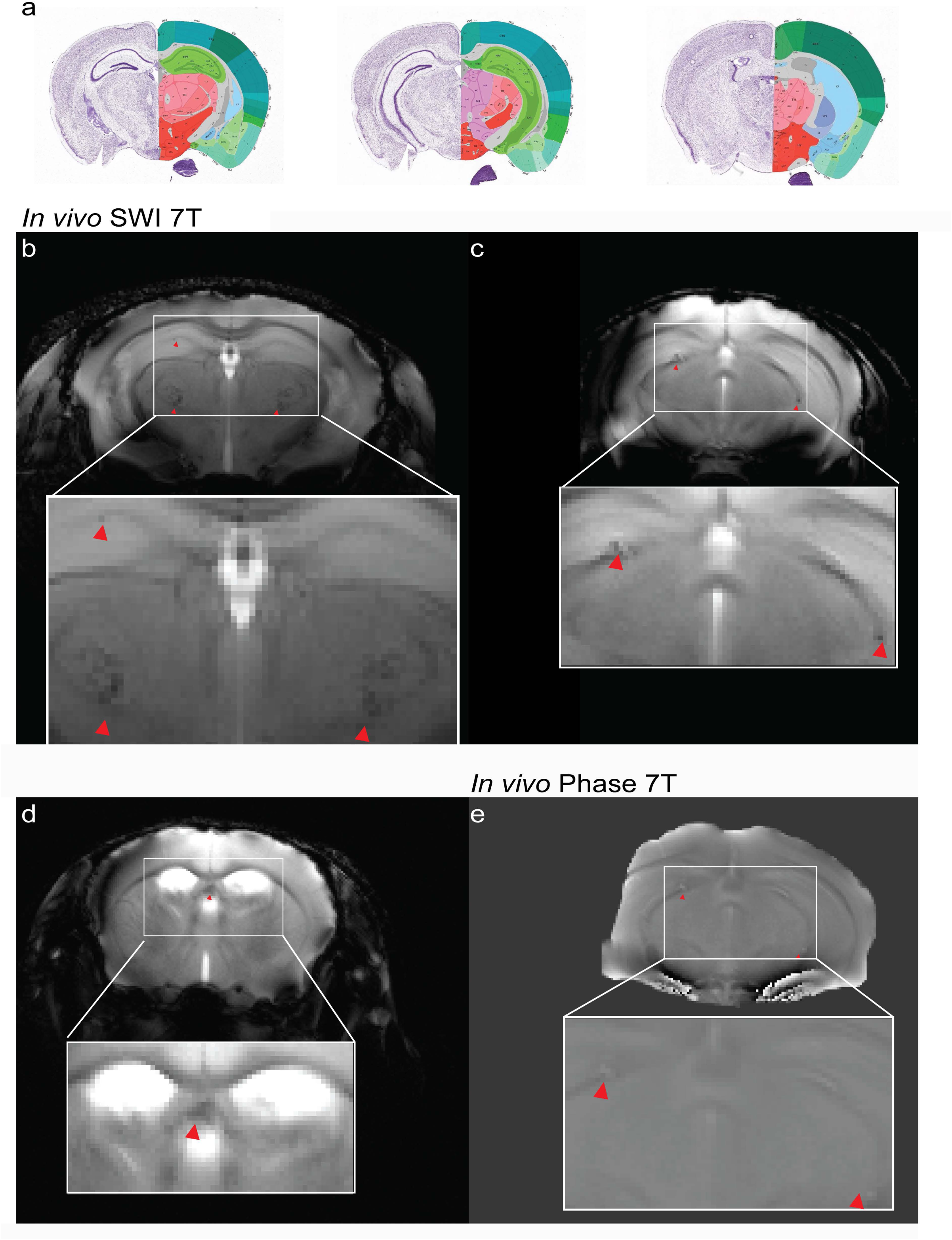
Susceptibility weighted/phase magnetic resonance imaging reveals characteristic pattern *in vivo* in P301L mouse brain. (a) Coronal sections from a mouse brain atlas [24] show location of susceptibility weighted (SW) and phase images; (b) Representative *in vivo* SW and (c) phase image at 7T showing hypointensities/positive phase shifts (red arrowheads) in the thalamus of an 18 month-old P301L mouse and (d) choroid plexus of the third ventricle of a 5 month-old P301L mouse; (c, e) SW and corresponding phase image in the hippocampus of a 5 month-old P301L mouse.

Hypointensities/positive phase shifts in P301L mice were detected in all age groups by using *in vivo* and *ex vivo* SWI/phase image, prominently in the hippocampus, but also in the thalamus, caudate nucleus, choroid plexus and midbrain (**Fig. 1, 2, 3**). In aged non-transgenic littermates such lesions were only observed in the choroid plexus (e.g. positive phase shifts in the 4^th^ ventricle, which appeared as a cohesive structure) (**Fig. 2a-c**). In the other brain regions hyperintenities/positive phase shifts on SW/phase images appeared as ovoid (**Fig. 3b, c, f, g**), elongated (**Fig. 3e, i**) lesions, and nests (**Fig. 3d, h**). D’Agostino & Person normality test results in p = 0.0648, indicating passing the normality test. The number of SW/phase diamagnetic deposits increased with age in the hippocampus, but not the other brain structures (18 months-old *vs* 3 months-old, p <0.0001; *vs* 5 months-old, p <0.0001; and *vs* 9 months-old, p <0.0001, **Fig. 3j**). No statistically significant difference was observed between male and female mice within age groups (**Fig. 3k**).

**Figure 2.**
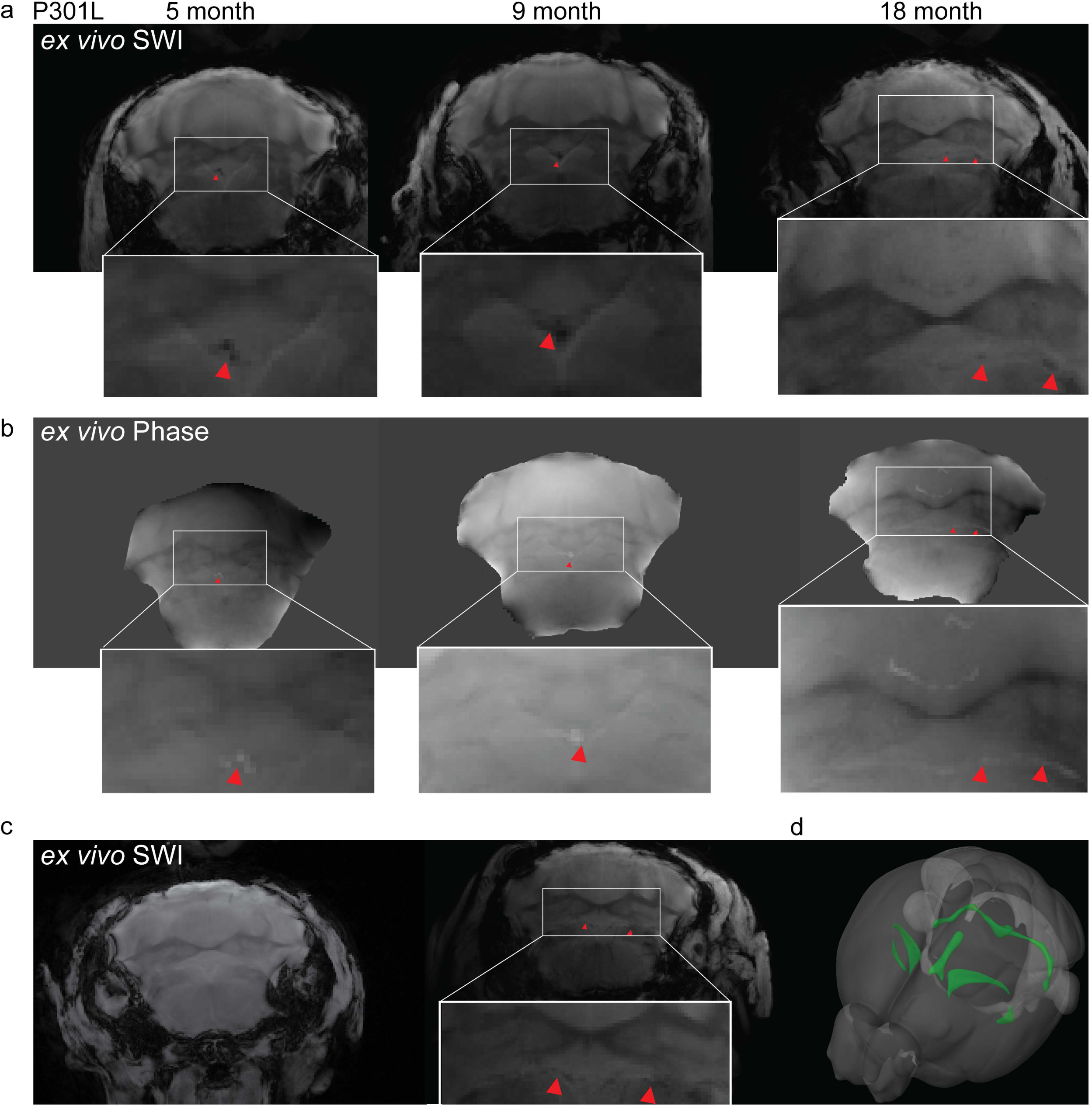
Comparison of imaging findings in the choroid plexus of P301L and non-transgenic mice. (a) Representative *ex vivo* susceptibility weighted (SW) magnetic resonance imaging at 9.4T and (b) corresponding phase image showing hyperintensities in the fourth ventricle choroid plexus of 5, 9 and 18 month-old P301L mouse (red arrowheads); (c) Representative *ex vivo* SW images at 9.4T in the fourth ventricle choroid plexus of 9 month-old and in 5 month-old non-transgenic littermates (NTL); (d) 3D mouse brain atlas from Allen Institute with choroid plexus highlighted in green [24].

**Figure 3.**
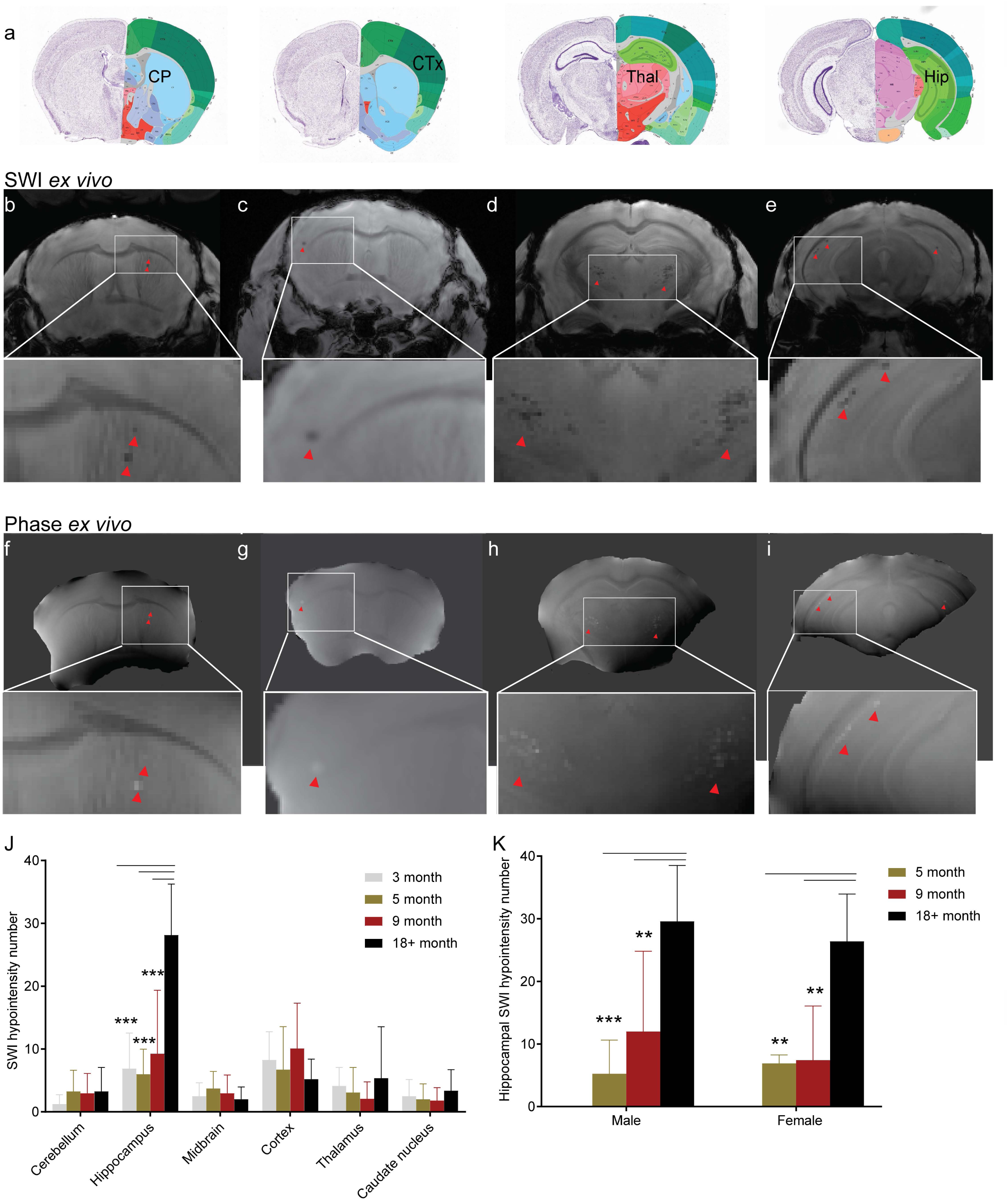
Magnetic resonance imaging reveals imaging pattern in *ex vivo* brain of P301L mice. (a) Coronal mouse brain atlas from Allen Institute (slices 46, 56, 76, 89) [24] showing corresponding location on magnetic resonance images; (b-i): Representative *ex vivo* susceptibility weighted (SW) and phase image at 9.4T showing (b, f) hypointensities/positive phase shifts in the 5 month-old P301L mouse; (c, g) in the cortex of a 9 month-old P301L mouse; (d, h) in the thalamus of a 18 month-old P301L mouse; (e, i) in the hippocampus of a 18 month-old P301L mouse (red arrowheads) (j) Quantification of regional distribution of SW hypointensities. 18+ month (n = 11) compared to 3 month (n = 4, p < 0.0001), 5 month (n = 11, p < 0.0001), 9 month (n = 10, p < 0.0001); (k) Number of SWI hypointensities in male and female P301L mice of all age groups; **p <0.01, *** p < 0.001 two-way ANOVA with Turkey’s *post hoc* analysis.

### µCT detects hyperdense lesions in the brain of P301L mice

To further assess cerebral lesions in the brain of P301L mice, *ex vivo* μCT scans were performed on the same brain samples. μCT images showed hyperdense lesions in 5, 9- and 18+ months-old P301L mice. The higher X-ray density in these areas indicates the presence of material of higher atomic number than soft tissue [56]. Hyperdense lesions were seen in regions that correspond to the hippocampus, deep brain regions, cerebellum, and cortex when compared to SW and phase MR images in the same mice (**Fig. 4a-d**). Hyperdense structures in the choroid plexus of the third ventricle were observed in the μCT images of P301L mouse brains and also in aged controls. However, fewer lesions were observed using μCT compared to MRI. Nevertheless, this adds further evidence for the presence of intracranial calcification in P301L mice.

**Figure 4.**
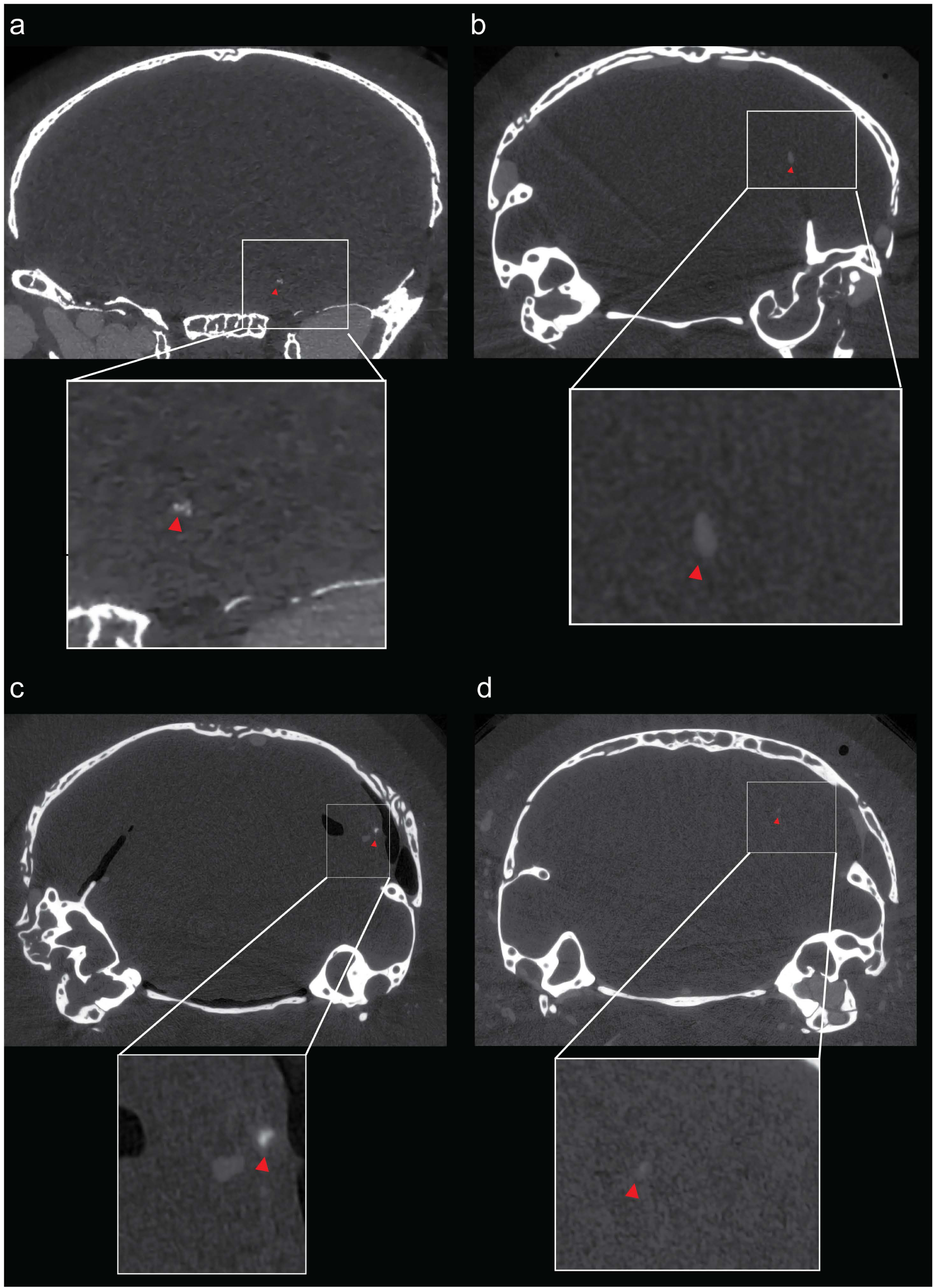
Hyperdense lesion detected on micro computed tomography images of the brains of P301L mice. (a-d) Representative *ex vivo* micro computed tomography showing hyperdense lesions in different brain regions (red arrowhead).

### Vascular bone protein-containing nodules in the thalamus of P301L mice

As a next step, we used histochemistry and immunohistochemistry to investigate the nature of the imaging lesions in different brain regions of P301L mice. To this end, we used immunohistochemical staining with antibodies against bone proteins deposited in brain calcifications [15, 75]. In the thalamus, osteocalcin- and osteopontin-positive nodules were associated with blood vessels, in the absence of phosphorylated-tau (AT8 and AT100) staining (**Fig. 5a-h**). Interestingly, similar to previous reports [42, 74], we detect amyloid-precursor protein (APP), a marker for damaged neurons [58], deposition in vascular nodules containing bone proteins (**Fig. 5i-l**). These nodules were visualized using hematoxylin and eosin staining, a standard histological stain, indicating the basophilic nature of these structures (**Fig. 5m**). In addition, vascular calcification in the thalamus elicits a strong glial reactivity (**Fig. 5n, o**), reminiscent to changes described in a mouse models of primary familial brain calcification [26, 42, 75].

**Figure 5.**
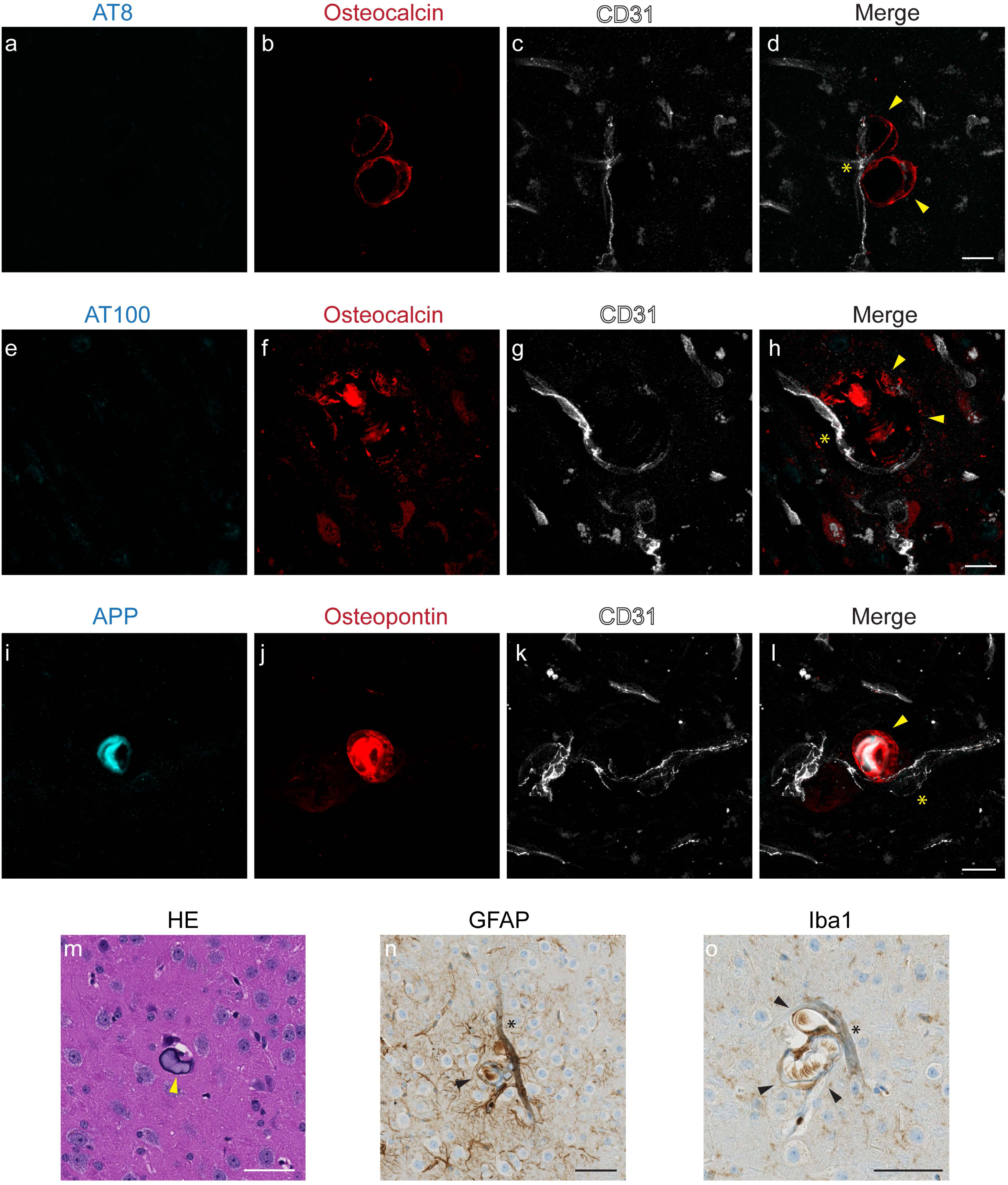
Characterization of vessel-associated nodules in the thalamic region of 18 month-old P301L mice. (a-l) Osteocalcin-positive (red) nodules (b, f, j) are associated with vessels (c, g, k, CD31; white) and do not stain with antibodies recognizing different phosphorylated residues of tau, AT8 (a) and AT100 (e); Thalamic nodules are positive for amyloid-precursor protein (i, APP, cyan) and osteopontin (j, red); (m) Hematoxylin and Eosin (HE) stain of thalamic nodules; (n, o) Thalamic nodules elicit glial reactivity. Activated astrocytes (n) and microglia (o) surround vascular nodules. Vessels are visualized using CD31 (c, g, k, white) and vessel adjacent to bone protein containing nodule is marked using an asterix (d, h, l). Arrowheads (d, h, l) mark thalamic nodules. Scale bars: 15 µm (a-l) and 50 µm (m-o).

We performed histology on sagittal brain sections of P301L mouse and non-transgenic littermate brains to verify the presence of other neuropathology in the brain of P301L mice. Alcian blue, Periodic acid–Schiff, and Hematoxylin & Eosin staining showed no pathological abnormalities or infarction in the mouse brain P301L mice at 18 month-of-age (**SFig. 1**). Furthermore, Prussian blue staining demonstrated that the imaging pattern in the P301L mouse brains were not cerebral microbleeds (**SFig. 1c, d**).

### Vascular and intracellular bone protein-containing nodules s in the hippocampus

The hippocampus of P301L mice was found to be a prominent site of imaging pattern indicative of tissue calcification (Fig. 3). Immunohistochemistry using antibodies against bone proteins confirmed their presence in deposits in the hippocampus (**Fig. 6a-p**), which were vascular (**Fig. 6e-h**) and parenchymal (**Fig. 6i-l**). In contrast to the thalamus, we observed in the hippocampus of P301L mice intracellular staining of osteocalcin which was co-localizing with phosphorylated-tau detected using antibodies AT8 and AT100 (**Fig. 6i-l**).

**Figure 6.**
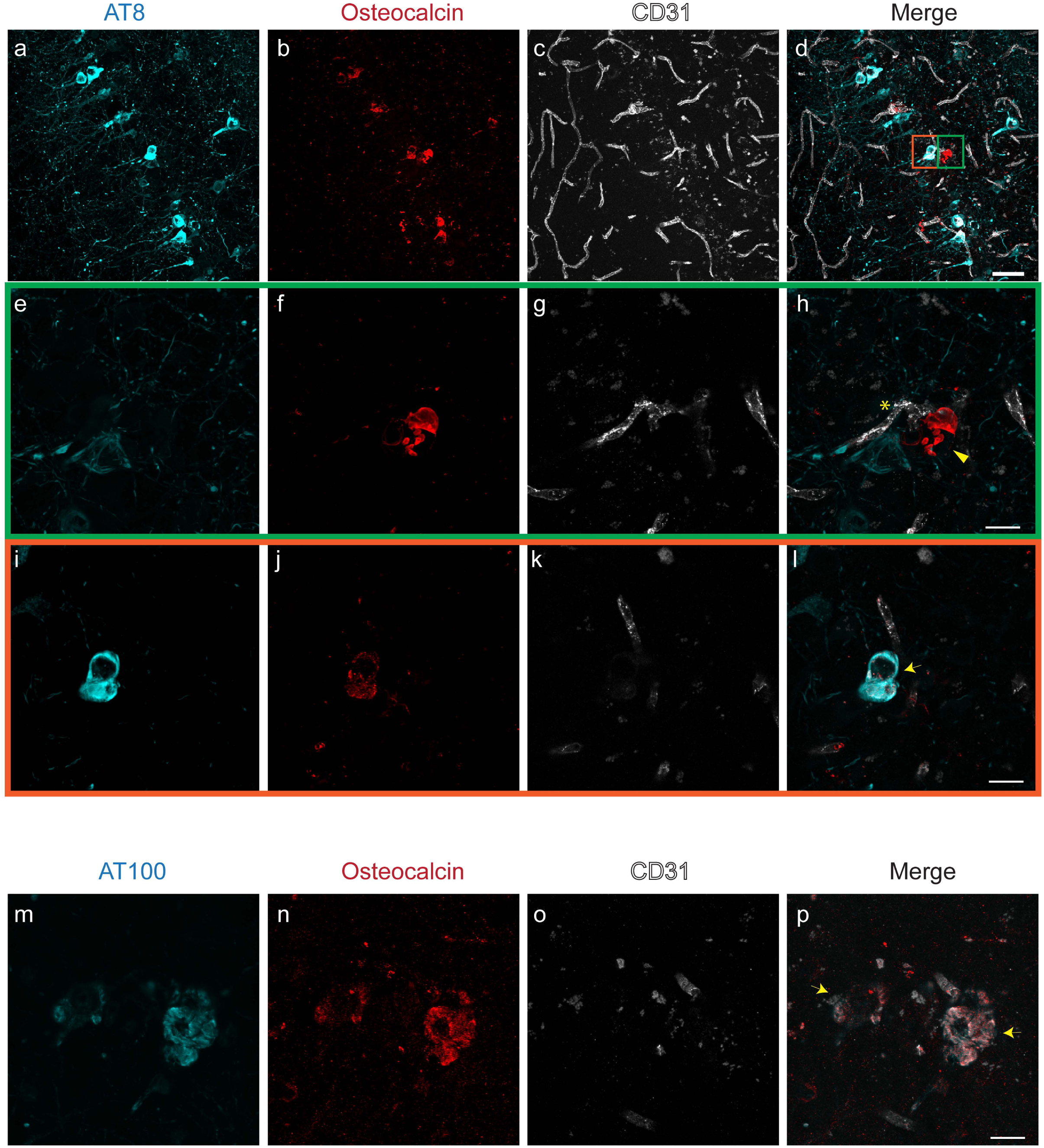
Characterization of hippocampal inclusions in 18 month-old P301L mice. (a-p) Osteocalcin (b, f, j, red) staining in the hippocampus. (e-h, green inset) Extracellular osteocalcin-positive (f, red) nodule is vessel-associated (g, CD31; white) but did not stain with the phosphorylated-tau AT8 (e, cyan) antibody. (i-l, orange inset) Intracellular co-localization of phosphorylated-tau AT8 (i, cyan) and osteocalcin (j, red) in a hippocampal neuron; (m-p) Co-localization of osteocalcin (n, red) positivity with phosphorylated-tau AT100 (m, cyan) in the hippocampus of P301L mice. Vessels are visualized using CD31 (c, g, k, o, white) and vessel adjacent to nodule is marked using an asterix (h). Arrowheads (h) mark extracellular nodules and arrows (l, p) mark intracellular co-localization of phosphorylated-tau stains and osteocalcin. Scale bars: 50 µm (a-d) and 15 µm (e-p).

## Discussion

In this study, we observed imaging pattern on high-resolution μCT and gradient recalled echo MRI, indicative of intracranial calcification, in brains of P301L mice. Immunohistochemistry revealed that these bone protein-containing lesions were either vessel-associated or intracellular and co-localizing with phosphorylated-tau.

CT is considered the non-invasive gold standard for the identification of intracranial calcification [3], with hyperdensity threshold >100 Hounsfield units [5]. In comparison, in conventional T_1_-, T_2_-weighted spin echo MR images calcifications appear with various signal intensities, which makes differentiation of calcified tissue from other sources of focal contrast (hemorrhagic lesions or infarctions) difficult. On the other hand, SW and phase imaging, two gradient recalled echo MRI techniques, allow distinguishing tissue calcification from cerebral microbleeds based on the magnetic susceptibility of the lesions [55]. In our study, all hypointense inclusions on SW images of P301L mouse brain were diamagnetic, indicating the presence of intracranial calcifications (**Fig. 1-3**). Negativity for Prussian blue staining indicated the absence of cerebral microbleeds and other iron deposition in the brain of P301L mice (**SFig. 1c, d**). Immunohistochemistry revealed that lesions are positive for bone proteins, thus indicating that the characteristic radiological pattern corresponds to markers of calcified tissue (**Fig. 5, 6**). Studies using clinical brain imaging data suggest that the diagnostic accuracy of SWI outperformed conventional MRI and CT in detecting brain calcifications [1, 32, 71, 72, 76], although the diagnostic accuracy of different methods might depend on disease indication, MRI parameters and field strengths used. In non-transgenic littermates, imaging abnormalities were found confined to the choroid plexus. While this has been reported to occur in aged individuals [20] it has so far not been reported as a phenotype in aged mice. In P301L mice we observed high inter-individual variations in hypointensities on SW images in all anatomical regions (**Fig. 2g**), similarly to previous studies in patients with familial brain calcification and mouse models thereof [26, 75].

The etiology of most forms of brain calcifications are unknown, but they may occur under several pathophysiological conditions, including inflammation, metabolic, infectious and genetic syndromes and after exposure to toxins or radiotherapy [10, 12, 35]. Loss-of-function mutations in genes involved in familial forms of diseases presenting with brain calcification have been associated with disturbance in phosphate homeostasis and the dysfunction of the blood-brain barrier [34]. This was supported by the finding of mutations in tight junction components in other genetic diseases with intracranial calcifications [40, 45]. Moreover, studies using hypomorphs of platelet derived growth factor subunit B suggested a connection between blood-brain barrier impairment and brain calcification [26], however, later studies did not find a causal evidence [65, 66]. Interestingly, blood-brain barrier breakdown has been reported in P301L tauopathy mice, with erythrocyte and leukocyte infiltration occurring before accumulation of hyperphosphorylated tau [4], thus it would be of interest for future studies to investigate if blood-brain barrier impairment plays a role in the manifestation of brain calcification in this setting.

The genetic cause of basal ganglia calcification with dementia and bone cysts has been linked with triggering receptor expressed on myeloid cells 2 (*TREM2*) mutation [28], and TYRO protein tyrosine kinase binding protein (TYROBP, formerly DAP12) [48] on chromosome 19. Mutations in TREM2 presenting as a FTD-like syndrome without bone involvement were later also reported [8, 17, 21, 22]. TREM2 deficiency exacerbates tau pathology through dysregulated kinase signaling in the tauopathy mouse model [2], suggesting microglial activation may also play a role in the tau pathology. We observed strong glial reactivity towards calcified nodules in the thalamus of the P301L mice (**Fig. 5n, o**). Reactive astrocytes and microglia surrounded calcified deposits and it will be interesting to study whether glia cells control the progression of the pathology as in hereditary brain calcification [74]. Further studies will need to elucidate if these calcifications in P301L are also TREM2 dependent and if there will be differences in susceptibility between the thalamic and hippocampal ones.

Intracranial calcification may also be linked to tauopathy. Studies demonstrated hippocampal calcification in the brain of patients with AD and FTD using CT and histopathology [51]. This is in line with our study where hypointensities were most abundant in the hippocampus, increasing in an age-related manner (**Fig. 3j**). Furthermore, we found co-localization of tau (AT8, AT100) with osteocalcin in hippocampal neurons, an earliest brain region affected in AD and FTD (**Fig. 6d, l**). In P301L mice, tau accumulation is prominent in the hippocampus, amygdala, and cortical regions, but sparse in the brain stem, thalamus, and caudate nucleus [18, 69]. The tau distribution was not matched regionally with the hypointensity pattern seen on MR imaging at whole brain level (**Fig 1, 2, 3**). The susceptibility to develop calcification may thus depend on a number of other factors in addition to human tau overexpression, such as the biochemical, and genetic properties. The individual and combined contribution of these factors to the formation of brain calcification needs further research [39]. Given the non-invasive nature of the imaging read-outs, longitudinal studies would be informative with respect to individual changes in calcium load and can be combined with other read-outs (e.g. perfusion measurement, elastography etc.).

We observed APP and osteopontin accumulation in the thalamic nodules in P301L mice. APP is a marker for damaged neurons [58] and osteopontin is a bone protein upregulated in damaged brain [59]. Both APP [58] and osteopontin [15, 75] are reported in the extracellular matrix of calcified tissue. The presence of APP could be an indication of damaged neurons in the area [42]. Here, we observed that they accumulate in the thalamic calcifications in P301L mice similar to other reports about brain calcifications, further supporting that these nodules are indeed calcified. As the antibody for APP binds to amino acids 85-99 of the C99 fragment of APP, which might bind to amyloid-beta deposits and APP C-terminal fragment C99 located in dystrophic neurites.

Once manifested, brain calcification can affect brain function by evoking expression of neurotoxic astrocyte markers, interfering with neuronal circuitry and/or glucose metabolism [32, 60, 75]. However, the functional contribution of brain calcification to neuropsychiatric symptoms in various brain disorders is still debated [13, 47]. Patients with primary familial brain calcification often present with impaired movement (parkinsonism and dystonia), but also cognitive impairment and psychiatric manifestations, including schizophrenia-like symptoms, mood disorders, or obsessive–compulsive disorder [13]. Patients with diffuse neurofibrillary tangles with calcification including early but progressive memory and verbal disturbances, followed by psychiatric symptoms. While the P301L strain shows impaired memory functions in hippocampus- and amygdala-dependent tasks [49, 50], future studies need to disentangle the potential contributions of tauopathy-related brain calcification to functional deficits.

## Conclusions

We described a new brain phenotype of P301L mice, characterized by imaging patterns of intracranial calcification. The study further suggests a potential link between tau deposition and tissue calcification, where the underlying pathophysiology and functional consequences need to be further investigated. The P301L mouse strain therefore may be a suitable model to study the pathogenesis and pathophysiology of brain calcifications in FTD and AD, and other tauopathies.

## Availability of data and material

The datasets generated and/or analyzed during the current study are available in the repository (DOI: 10.5281/zenodo.3518986).

## Author Contributions

RN, AK, JK conceived and designed the study; RN, YZ, YY, GK performed the experiments; RN, YZ, GK, LK, AK and JK interpreted the results; RN and JK wrote the manuscript; all coauthors contributed constructively to the manuscript.

## Acknowledgement

The authors acknowledge Prof. Daniel Razansky and Prof. Markus Rudin at the Institute for Biomedical Engineering, ETH Zurich & University of Zurich for access to the infrastructure of the animal imaging facility; Dr. Zsofia Kovacs, Dr. Mark-Aurel Augath at the Institute for Biomedical Engineering, ETH Zurich & University of Zurich, Daniel Schuppli at the Institute for Regenerative Medicine, University of Zurich and Dr. Gabriella Bodizs, ScopeM, ETH Zurich for technical support. Dr. Joanne Lim for language corrections.

---

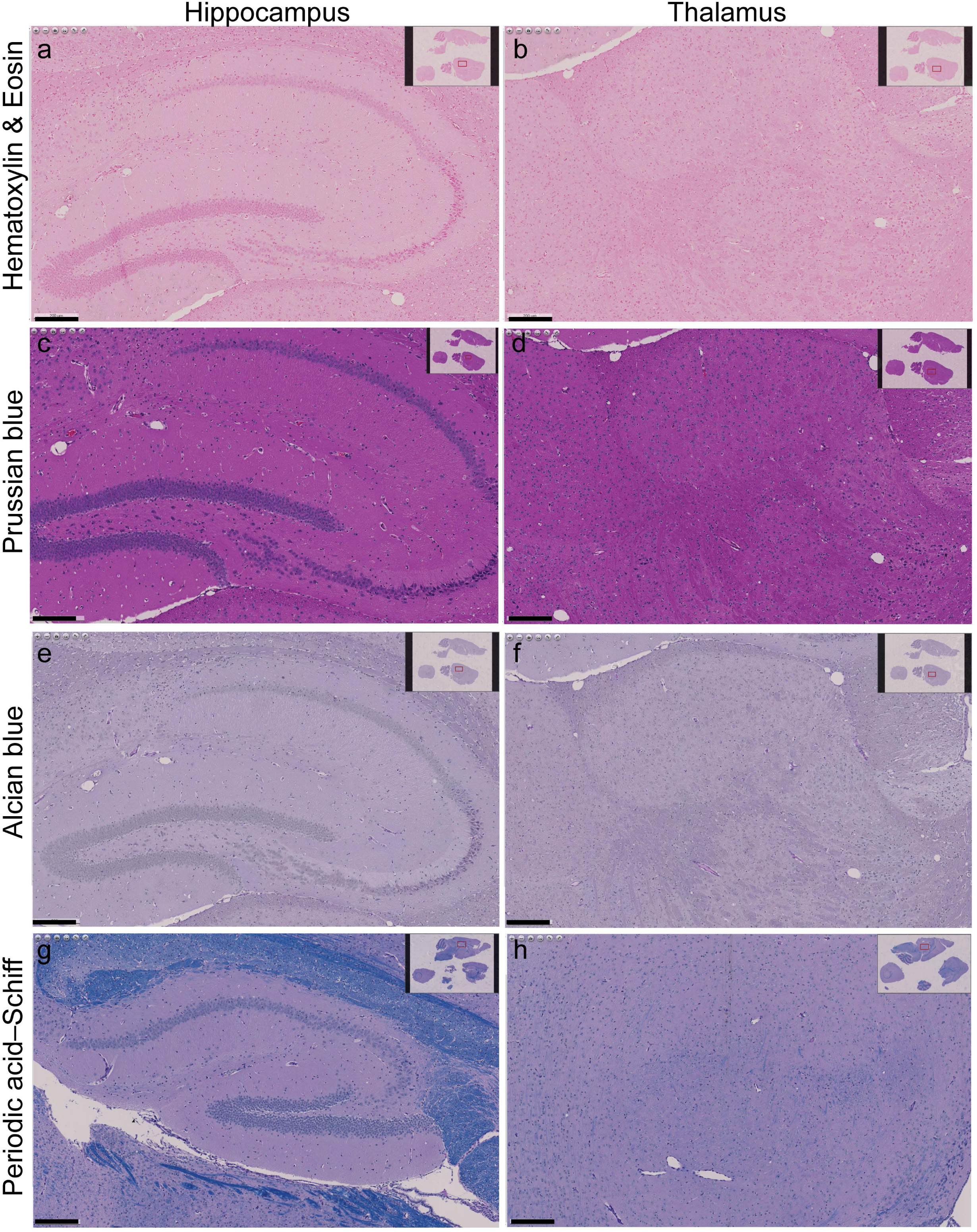

