## supplemental table and figure legend for "Tau deposition is associated with imaging patterns of tissue calcification in the P301L mouse model of human tauopathy"

**Supplementary table 1 Summary of transgenic and non-transgenic mice used in the study**

|  | Homozygote (P301L) | Non-transgenic |
| --- | --- | --- |
| 3 month | 4M |  |
| 5 month | 5F/6M | 5F/1M |
| 9 month | 6F/4M | 2F/7M |
| 18+ month | 5F/6M |  |

M: male; F: female; Homozygote and non-transgenic mice have the C57B6 background.

**Supplementary table 2. List of primary antibodies used for immunohistochemistry**

| **Antibody** | **Company** | **Cat. No.** | **Dilution** |
| --- | --- | --- | --- |
| Goat anti-Osteocalcin | Alfa Aesar | J65216 | 1:500 |
| Rabbit anti-APP | ThermoFischerScientific | PA5-19923 | 1:200 |
| Mouse anti-AT100 | ThermoFischerScientific | AB223652 | 1:100 |
| Mouse anti-AT8 | ThermoFischerScientific | AB223647 | 1:100 |
| Goat anti-Osteopontin | R&D systems | AF808 | 1:100 |
| Rat anti-CD31 | Dianova | DIA-310 | 1:100 |

**Supplementary Fig. 1 Histology staining in the hippocampus and thalamus of representative 18 month-old P301L mice (**a, b) Hematoxylin & Eosin; (c, d) Prussian blue; (e, f) Alcian blue; (g, h) Periodic acid–Schiff. Scale bar 200μm.
